# A 3D printed plastic frame deeply impacts yeast cell growth

**DOI:** 10.1101/2023.06.16.545257

**Authors:** Esther Molina-Menor, Àngela Vidal-Verdú, Carlos Gomis-Olcina, Juli Peretó, Manuel Porcar

## Abstract

Solid State Fermentation (SSF) processes have been explored for yeast growth and protein and metabolites production. However, most of these processes lack standardization. In this work, we present a polylactic acid (PLA) 3D printed matrix that dramatically enhances yeast growth when embedded in liquid media compared to equivalent static cultures, and changes yeast expression patterns at the proteome level. Moreover, differences in sugar assimilation and ethanol production, as the main product of alcoholic fermentation, are observed. Our results suggest that these matrixes may be useful for a vast range of biotechnological applications based on yeast fermentation.

## Introduction

Microorganism-culturing techniques have drastically evolved in the last years. Specifically, since the development of high-throughput DNA sequencing technologies, several groups have worked in the improvement of culturing conditions in order to increase the number of microbial species grown in labs and to improve their growth (Overmann et al., 2017).

Industrial microorganisms are mainly grown in planktonic, continuous (chemostat) or batch cultures (Carsanba et al., 2021; Jiao et al., 2022). In particular, yeast industrial production can be approached through different bioreactor configurations, but all of them usually in liquid or submerged fermentation (SmF). On one hand, batch culturing is the simplest and cheapest strategy, not only from the economic point of view but also considering the workload. In batch culturing, the culture is prepared once, simplifying sterilization and stock preparation. However, the main disadvantage is growth inhibition due to the accumulation of metabolic products, some of which are toxic (Carsanba et al., 2021). On the other hand, continuous cultures avoid this limitation by uninterrupted feeding and removing of products. It allows monitoring and comparing in real time different parameters, but it is not always a feasible strategy as some products may require the culture to reach stationary phase to be synthesized (Carsanba et al., 2021). In between there can be found fed-batch culturing, the most widely used strategy, in which the effect of growth inhibition is lower than in conventional batch (Carsanba et al., 2021).

Mimicking the environmental conditions that microbes find in their natural habitats is one of the aims of culturomics, not only for cultivating more species but also to improve their growth in terms of production performance, and also in terms of energy and economic savings. For example, some authors studied the effects on cell immobilization over the conventional batch mode in the synthesis of specific compounds (citronellol) and found a lower subproduct production (Esmaeili et al., 2012), improving, thus, the recovering of the product. For their part, Raghavendran et al. (2020) developed a system to reduce the cost of aeration of batch yeast cultures by using a fluidic oscillator that generated microbubbles. Moreover, some authors have worked on the growth of bacteria on solid surfaces in order to enable biofilm formation, as some bacterial species require this for optimal growth (Gich et al., 2012).

Solid-State Fermentation (SSF) has been explored and partially developed as a culture technique that improves oxygen transfer, avoids foaming in cultures and mimics the natural growth conditions of some microorganisms, such as filamentous fungi, yeast or biofilm-forming bacteria (Hölker and Lenz, 2005; Lima-Pérez et al., 2019). This microbial-growth alternative is based on the fermentation of microorganisms on an organic solid moist surface, such as agricultural waste, or inert solid surfaces like polyurethane foam (PUF) to which nutrients are added. Although there are some studies on the promising role of SSF in the biosynthesis of specific compounds and the advantages that it holds over SmF, at industrial scale, the growth of microorganisms to produce either biomass or secondary metabolites is still mainly SmF (López-Pérez and Viniegra-González, 2016; Singhania et al., 2009; Valdés-Velasco et al., 2022).

In the line of this, here we report an attempt to standardize yeast growth on solid matrixes with an unprecedented approach. Based on a preliminary test on sand, cotton and sponges, we describe a culturing strategy consisting on geometrical 3D solid tough polylactic acid (PLA) matrixes embedded in liquid media for sourdough yeast *Saccharomyces cerevisiae* production and analyze how culture conditions modulate their growth and their proteomes.

## Materials and Methods

### 3D-matrixes design

3D-printed cylinders were designed by Marker Station 3D (Valencia, Spain). The design consisted in a cylindric spiral with regular holes (1X scale: 23.7 × 24.4 × 94.8 mm). The designs were printed in black tough polylactic acid (PLA) in an Ultimaker S3 3D printer (Figure 1).

**Figure 1.**
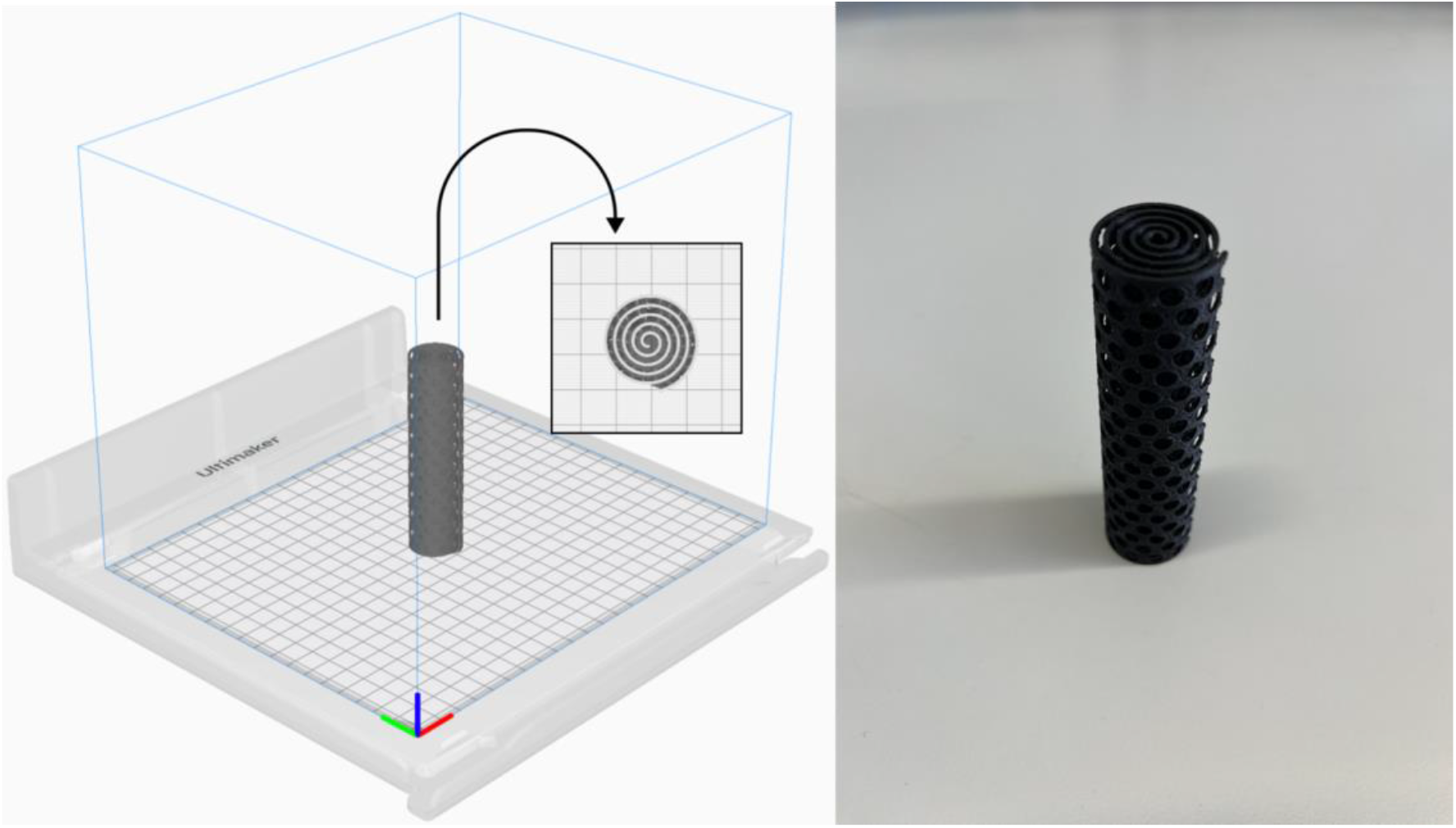
3D printed PLA matrix design.

### SSF experiment

Yeast precultures were grown overnight in YPD medium (in g/L: 10 yeast extract, 20 peptone, 20 dextrose) at 30 °C. Cultures were adjusted to a final optical density of 600 nm (OD_600_) of 0.1 in YPD. SSF matrixes were submerged in 1/10 (v:v) bleach in order to disinfect them for 20-30 min. Then, they were thoroughly washed in sterile distilled water in three cleaning steps. The SSF matrixes were placed in 50 mL tubes, and 25 mL of the adjusted yeast culture was poured into them. The growth controls consisted of 25 mL of culture in 100 mL flasks for agitated/shaking conditions (SK) (180 rpm) and 25 mL of culture in 50 mL tubes for static conditions (ST). The experiments were carried out in triplicate (biological replicates). A tube without yeast was used for sterility control. All tubes were incubated overnight at 30 °C and centrifuged in 50 mL tubes for 5 min at 4500 rpm. After discarding the supernatant, cells were resuspended in sterile PBS for cleaning. A centrifuging step was repeated under the same conditions. After discarding the supernatant, cells were finally resuspended in sterile PBS at a final volume of 25 mL. OD_600_ was, then, measured. One-way ANOVA followed by Turkey’s multiple comparisons test was performed using GraphPad Prism version 9.0.0 for macOS, GraphPad Software, San Diego, California USA, www.graphpad.com.

### Metabolite measurement

D-glucose and ethanol content were measured in overnight culture supernatants after centrifuging the cultures for 10 min at 5000 rpm in a Y15 Automated Analyzer (BioSystems, Barcelona, Spain). One-way ANOVA followed by Turkey’s multiple comparisons test was performed using GraphPad Prism version 9.0.0 for macOS, GraphPad Software, San Diego, California USA, www.graphpad.com.

### Proteomics

One experiment (in triplicate) was carried out for proteome analysis. After collecting and cleaning the cells, pellets were directly frozen and stored at -80°C until required. Proteomic analysis was carried out by the Central Service for Experimental Research (SCSIE, Universitat de València). Proteins were separated by 1D SDS PAGE electrophoresis and identified through LC-MSMS. Differential proteomic analysis was carried out through SWATH. Proteomic analysis was done by Darwin Bioprospecting Excellence SL (Paterna, Valencia).

## Results

### SSF experiment

Preliminary assays of SSF approaches using solid substrates such as sand, cotton and sponges revealed enhanced yeast growth (data not shown) compared to traditional SmF cultures. Therefore, an experiment with standardized 3D matrixes was designed. Growing yeast in alternative culture approaches (liquid shaken cultures-SK, liquid static cultures-ST and 3D-SSF cultures) resulted in significantly different cell yields measured by OD (Figure 2A) (p-values<0.05). Specifically, the highest biomass was obtained through conventional SK cultures, followed by 3D-SSF and ST culturing. Although 3D-SSF culturing did not result in similar cell-yields compared to SK, it significantly enhanced growth compared to the equivalent submerged culture without agitation (ST).

**Figure 2.**
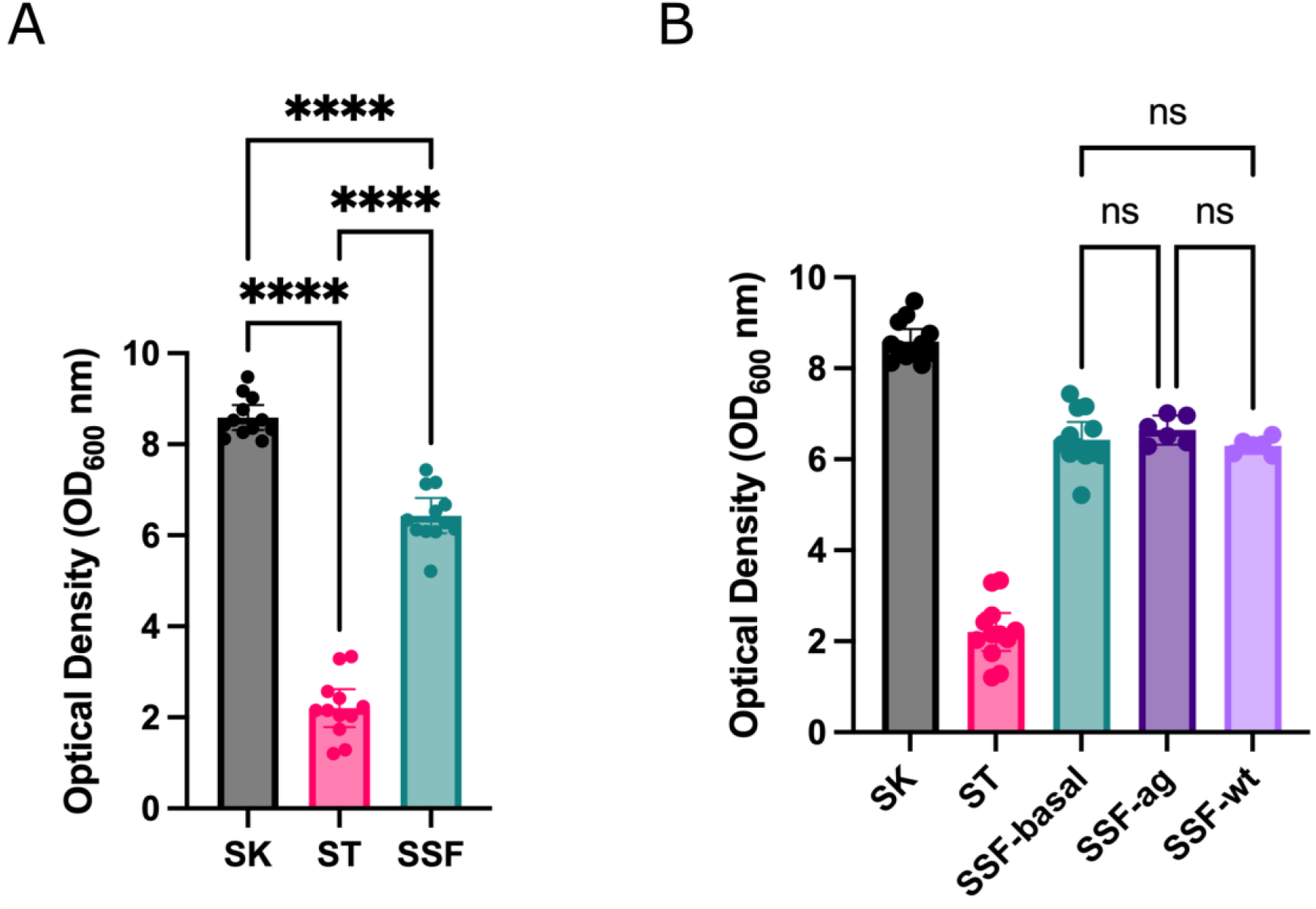
Yeast growth (OD_600_ nm) inside 3D matrixes. Shaking and static controls are included. Confidence interval of twelve replicates (biological replicates) is represented in the error bar. (A) SSF experiment. (B) SSF experiment in different configurations: with the PLA matrix and static conditions (3D basal), agitated to increase bubbles or hit to remove any air bubble (without bubbles; wt). Shaking and static controls are included.

In order to determine whether the observed increased biomass may result from increased oxygen through embedded bubbles inside 3D matrixes, an experiment was carried out in which tubes with 3D matrixes were subjected to agitation prior to incubation or hit in order to remove air bubbles. However, no significant differences were found among these three conditions and the rest of the controls displayed values accordingly to the previous experiments (p-values>0.05) (Figure 2B).

### Proteomics

Proteomic analysis was carried out in order to determine whether the observed enhanced growth in 3D matrixes was accompanied by differences in expression at the proteome level. The three conditions (agitated/shaking control (A), static control (E) and 3D-SSF matrix (N)) were analyzed in triplicate. Significant differences were found at the protein profile level as revealed in the PCA, in which there can be observed a clear separation of samples among conditions, as well as grouping of the biological replicates (Figure 3). Moreover, the number of differentially expressed proteins in the comparison of 3D-SSF with SK and ST controls was higher in the former than in the latter, suggesting that in terms of protein expression 3D-SSF culturing resembled more ST than SK fermentation.

**Figure 3.**
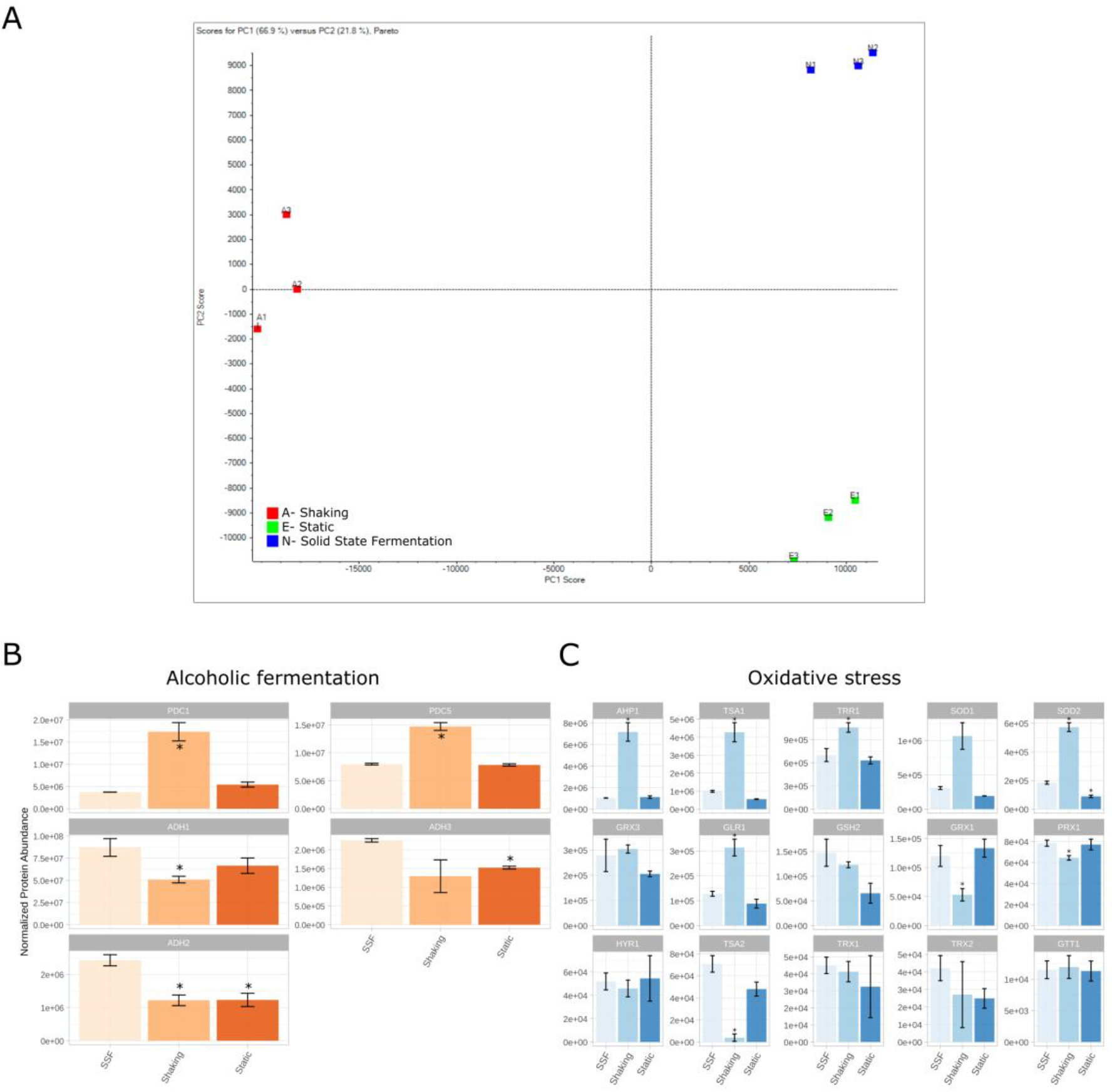
Proteomic analysis. (A) PCA showing the differential proteomic analysis (SWATH) of yeast cultures grown in the three conditions tested: agitated/shaking control (A), static control (E) and 3D matrixes (N). (B) Alcoholic fermentation protein expression levels. (C) Oxidative stress protein expression levels. * represents differential expression compared to SSF conditions.

With the purpose of determining the differences at the proteome level underlying this phenomenon, an analysis of differential expression of all the identified proteins was carried out. The proteins that were significantly over or under-represented in N-SSF samples with respect to A-shaking and E-static were identified prior to perform an enrichment analysis with STRING (https://string-db.org/), which is based on a protein-protein interaction database. This analysis allowed the identification of the metabolic pathways (KEGG pathways, https://www.kegg.jp/) that outstood in each of the groups with respect to the others. We hypothesized that metabolic routes involved in processes such as alcoholic fermentation and oxidative stress may display different expression patterns among conditions. Thus, a specific research and analysis on genes within these routes were performed (Figure 3B-C).

The analysis of proteins in SK and ST conditions compared to 3D-SSF revealed some patterns and significantly different expression levels. Regarding alcoholic fermentation proteins, the detected pyruvate decarboxylases (PDC1 and PDC5) were significantly more expressed in SK, while there were no differences for ST. PDC6 was not detected. In contrast, alcohol dehydrogenases (ADH1, ADH2 and ADH3) were in general terms overexpressed in 3D-SSF cultures with respect to SK conditions (Figure 3B).

Regarding proteins involved in oxidative stress, superoxide dismutases (SOD1 and SOD2), thioredoxin peroxidases (TSA1 and AHP1), glutathione reductase (GLR1) and thioredoxin reductase (TRR1) were overexpressed in SK cultures, in contrast to thioredoxin peroxidases TSA2 and PRX1, which were underexpressed in SK cultures (Figure 3C). SOD2 was also underexpressed in ST cultures, whereas the rest of the studied proteins (glutathione peroxidase (HYR1), glutathione transferase (GTT1), glutaredoxins (GRX1 and GRX3), glutathione synthase (GSH2) and thioredoxins (TRX1 and TRX2)) did not reveal clear differences among conditions.

### Metabolite measurement

Given the observed differences found at biomass yield and protein expression levels, we decided to explore the impact of fermentation in the cultures. Thus, D-glucose and ethanol contents were measured in the media after yeast growth and compared to the initial content in YPD medium (Figure 4).

**Figure 4.**
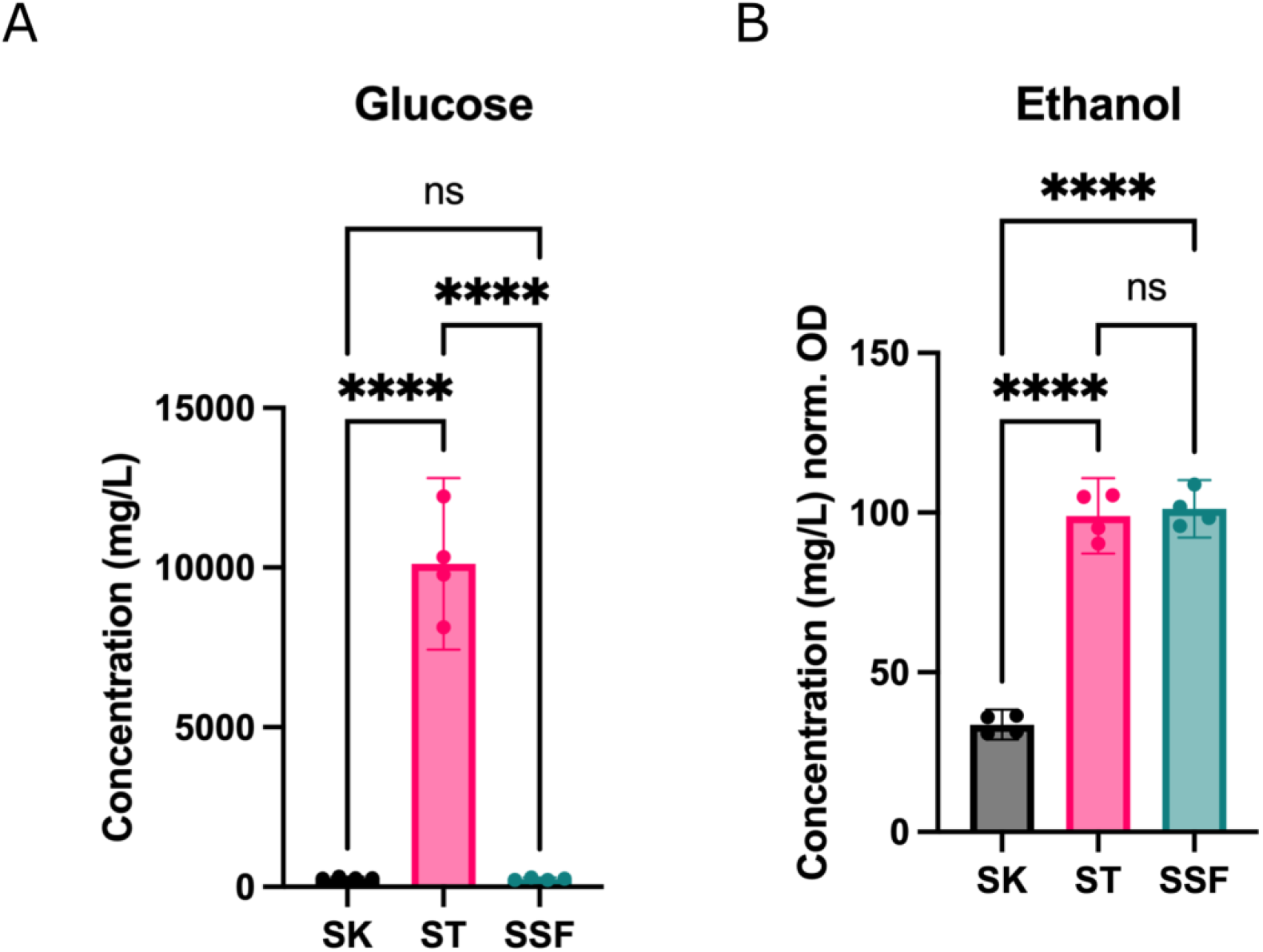
Metabolite profiles of cultures. (A) Sugar content (D-glucose) after fermentation (mg/L). (B) Ethanol production during fermentation (mg/L) normalized by cell density (OD_600_ nm).

YPD media was prepared at 20 g/L of D-glucose (quantified 19.32 g/L after autoclaving). D-glucose content was reduced to half in ST cultures (10.12 g/L), indicating poor sugar assimilation, whereas in both SK and SSF remaining sugar content was very low (280 and 245 mg/L, respectively) (Figure 4A). However, ethanol yields (normalized by the OD_600_ nm values achieved by each culture) were very similar in both ST and SSF cultures, and higher with respect to SK conditions (Figure 4B).

## Discussion

In this work we aimed at assessing whether the use of alternative culturing configurations based on standardized SSF inert matrixes may have an impact on yeast cell growth, not only in terms of the achieved cell densities but also in terms of protein expression patterns and metabolism. In the light of the results obtained, we conclude that solid, inert matrixes such as plastic cylinder results in a dramatic increase in yeast biomass production, and that different culturing setups result in different cell yields and also different protein profiles, which may be interesting from the biotechnological point of view and industrial biotechnology.

First, comparing yeast culturing under agitated and non-agitated (static) liquid cultures revealed, as expected, lower cell densities for static cultures. Microbial liquid cultures of aerobic organisms are usually agitated in order to favor the access to oxygen and nutrients, as oxygen poorly diffuses into water (Somerville and Proctor, 2013). Thus, it is expected that cultures incubated under shaking conditions achieve higher biomass yield. However, our results revealed that SSF matrixes embedded in liquid media and incubated under static conditions were able to enhance yeast growth with respect to the equivalent control culture in static (Figure 2).

We hypothesized, then, that 3D matrixes may trap oxygen molecules in bubbles that enhanced somehow growth. In fact, the use of inert media such as PUF increases the ratio area/volume favoring air spread and oxygen transfer in SSF (López-Pérez and Viniegra-González, 2016). Moreover, some authors have studied the effect of recipients in gas permeability and their impact on fermentation processes, such as the case of kimchi in onggi vessels, facilitating CO_2_ dispersion (Kim and Hu, 2023). However, no differences have been found when trying to, manually, change this condition (Figure 2B).

We explored how proteomes were affected by different incubation formats. The analysis of differential expression revealed clearly different protein profiles, and SSF cultures resemble more those of ST rather than SK ones, according to the distribution on the PCA and the number of differentially expressed proteins (Figure 3). As we hypothesized that both ST and SSF conditions (thus static ones) may favor alcoholic fermentation over respiratory metabolism, proteins involved in that pathway were specifically analyzed. In this regard, the overexpression pattern found for ADH1, ADH2 and ADH3 under SSF conditions, as well as to some extent under ST conditions, is in line with this hypothesis indicating enhanced fermentation in those cultures. Moreover, the profiles obtained for pyruvate decarboxylases (PDC1 and PDC5) are also coherent with their role in the cytosolic synthesis of acetyl-CoA required for lysine and lipid biosynthesis, essential processes for cell growth (Flikweert et al., 1996; Pronk et al., 1996).

Moreover, as a consequence of aerobic metabolism, Reactive Oxygen Species (ROS) and Oxidative Stress (OS) are generated. The analysis of the main proteins involved in the protection against OS and detoxification of ROS (see Herrero et al. 2008 for a review), revealed differences among culturing conditions. Specifically, the main defenses against OS associated to the operation of Krebs cycle (SOD1, SOD2, TRR1 and AHP1) were overexpressed under shaking conditions with respect to SSF. This is expected as respiratory metabolism is favored over fermentative pathways in the presence of oxygen. Moreover, the expression patterns found for TSA1 and TSA2 peroxiredoxins are also consistent with the literature. Both peroxiredoxins are homologue proteins that can complement one to the other, but TSA1 basal expression levels are significantly higher than those of TSA2 (Wong et al., 2002). In the case of SK conditions, TSA1 is enough to maintain its function, but in SSF and ST a complementation between TSA1 and TSA2 is observed (Figure 3C).

We also assessed sugar consumption and ethanol production (Figure 4), as D-glucose consumption is a common parameter measured to estimate the efficiency of SSF processes. For example, Christen et al. (1993) evidenced enhanced growth and total D-glucose consumption of *Candida utilis* grown on sugarcane. We expected higher use of sugars in agitated cultures given the higher OD observed, as well as higher ethanol biosynthesis in both static and SSF. D-glucose was almost completely used by yeast in shaking and SSF cultures (more than 99%), but around 50 % remained in static tubes (Figure 4A). However, ethanol yields normalized by OD in static and SSF were equivalent (Figure 4B). As this is unexpected, we hypothesize that under conditions of sugar depletion yeast may have started to consume ethanol, which could be supported by ADH2 overexpression, as it converts ethanol to acetaldehyde once preferable carbon sources are scarce and metabolism switches towards assimilation of nonfermentable carbon sources (Figure 3B).

As reviewed by López-Pérez and Viniegra-González (2016), SSF culturing strategies have been used for the production of proteins and metabolites by different yeast species, such as *Pichia pastoris, Kluyveromyces marxianus, K. lactis, S. cerevisiae* or some *Candida* spp., among others. The products obtained include proteins such as lipases and lytic enzymes, but also ethanol and aromatic compounds. Valdés-Velasco et al. (2022) described differences in the biosynthesis of lipopeptide biosurfactant in *Bacillus* strains being cultured under SmF or SSF, revealing that the use of PUF as an inert support for bacteria yielded higher productivity. Moreover, as reviewed by Wang et al. (2019) and described by Lahiri et al. (2021), incubating bacteria in static or in agitated conditions influences the biosynthesis of cellulose in terms of yield and form (either reticular films on the surface in static or sphere-like particles under shaking conditions) and offers different applications. Moreover, industrial biosynthetic processes may also consider the combination of different fermentation configurations, as Piecha et al. (2022) evaluated the production of polyhydroxyalcanoates (PHAs) and concluded that the combination of SSF and SmF would increase the productivity of the process. Altogether, alternative culturing strategies have an impact on the biosynthetic abilities of microorganisms and can be exploited for biotechnological benefit.

Although SmF has advantages for industrial processes, SSF holds potential for sustainable industrial production processes (Carsanba et al., 2021; Doriya et al., 2016). SSF has demonstrated to allow production with lower water requirement, which is of special interest in yeast as they naturally grow in low water availability environments, but also for bacteria that could adapt to these conditions (Couto and Sanromán, 2006; López-Perez and Viniegra-González, 2015). Moreover, SSF is an opportunity for recycling and revaluation of agricultural waste (Chilakamarry et al., 2022). That said, and to the best of our knowledge, inert, plastic matrixes have not yet been massively implemented to boost yeast biomass production.

Our studies serve as a starting point in the development of 3D structures that may change culture dynamics in terms of achieved biomass and metabolic fluxes for yeast-led industrial processes. Our findings arise some questions regarding the processes involved in the observed differences at biomass and proteome levels, and the metabolic fluxes. Further research is needed in order to elucidate the mechanisms underlying these results. In this regard, not only the development of matrixes with different geometric patterns or materials (e.g., autoclavable materials) would be of interest, but also the exploration of the role of other parameters that allow a deeper comprehension of the process are worthwhile studies. For example, sugar and ethanol quantification at different incubation times would give information about D-glucose consumption rates by each culture, ethanol synthesis rates and, in case of need, ethanol assimilation. Moreover, CO_2_ measurements may inform about the fermentation process. Also, investigating proteins related to functions such as quorum sensing or biofilm formation may shed some light on other processes that affect microbial growth.

Finally, many authors have outlined the limitations of SSF in large scale production processes (Chilakamarry et al., 2022; Piecha et al., 2022). Taken together, the results we show in the present article may be a first step towards a new view of SSF-like fermentation in yeast.

## Author Contributions

MP conceived the work. EMM, ÀVV and CGO performed the experiments. All the authors (MP, EMM, ÀVV, CGO and JP) analyzed the results, wrote and approved the manuscript.

## Acknowledgments

We are thankful to MarÍa Jesús Clemente and Felipe Werle for their help. We also thank Adriel Latorre, Daniel Torrent and Carmen Sanz from Darwin Bioprospecting Excellence S.L for their assistance with proteomic and metabolite analysis.

